# Towards comprehensive plasma proteomics by orthogonal protease digestion

**DOI:** 10.1101/2021.04.28.441706

**Authors:** Andrea Fossati, Alicia L. Richards, Kuei-Ho Chen, Devan Jaganath, Adithya Cattamanchi, Joel D. Ernst, Danielle L. Swaney

## Abstract

Rapid and consistent protein identification across large clinical cohorts is an important goal for clinical proteomics. With the development of data-independent technologies (DIA/SWATH-MS), it is now possible to analyze hundreds of samples with great reproducibility and quantitative accuracy. However, this technology benefits from empirically derived spectral libraries that define the detectable set of peptides and proteins. Here we apply a simple and accessible tip-based workflow for the generation of spectral libraries to provide a comprehensive overview on the plasma proteome in individuals with and without active tuberculosis (TB). To boost protein coverage, we utilized non-conventional proteases such as GluC and AspN together with the gold standard trypsin, identifying more than 30,000 peptides mapping to 3,309 proteins. Application of this library to quantify plasma proteome differences in TB infection recovered more than 400 proteins in 50 minutes of MS-acquisition, including diagnostic *Mycobacterium tuberculosis* (Mtb) proteins that have previously been detectable primarily by antibody-based assays and intracellular proteins not previously described to be in plasma.

## Introduction

Mass spectrometry-based proteomics is among the most promising technologies for biomarker discovery due to the ability to simultaneously detect thousands of proteins, post-translational modifications, and isoforms, all of which holds great potential as future biomarkers.^1^ This high throughput approach can lead to the identification of proteins that can be translated into simple, affordable, and non-invasive assays at the point-of-care for disease diagnosis and monitoring. For example, tuberculosis (TB) is a leading cause of mortality from an infectious disease globally for which diagnosis remains a key challenge. There is a critical need for rapid, low-cost, point-of-care assays but there are few promising biomarker ^2^ targets for assay development. Proteomics offers the potential to address this challenge and facilitate advances in diagnostics development for TB and other diseases.

Plasma is easy to obtain and has been used for diagnosis of a variety of infectious diseases, such as AIDS,^3^ Hepatitis C^4^ and recently, Sars-CoV-2. ^5^ The plasma proteome also represents a particularly challenging matrix to analyze due to the large dynamic range of protein concentrations spanning 10 orders of magnitude and the overwhelming presence of a select set of highly abundant proteins (e.g. albumin). Historically, this has limited both the number of proteins detected, as well as the reproducibility of detection. To mitigate issues in protein detection, numerous studies have successfully employed extensive off-line chromatographic fractionation, allowing for the injection of individual fractions of reduced complexity into the mass spectrometer. This approach has been highly successful to increase the number of proteins detectable in plasma, ^6,7^ albeit at the cost of a correspondingly dramatic increase in MS acquisition time to analyze dozens of fractions. Furthermore, the reliance on off-line fractionation introduces a low-throughput and cumbersome additional step in sample preparation that is not accessible to many labs.

Reproducible protein quantification is also critical for biomarker discovery, as differences in the abundance of specific proteins can be used as a clinical marker. Regardless of the method of quantification employed, data-dependent acquisition (DDA)^8^ strategies suffer from stochastic precursor ion sampling resulting incomplete quantification, particularly with increase sample numbers.^9^ In contrast, data independent acquisition mass spectrometry approaches (DIA/SWATH-MS) ^10^ sequentially sweep across m/z precursor isolation windows to acquire multiplexed tandem mass spectra irrespective of which peptides are being sampled. This results in highly complete and consistent quantification that readily scales for the analysis of hundreds or thousands of samples. While DIA offers great potential for plasma proteomics, most studies have been limited to measuring ≈ 300 proteins,^11^ partially due to the lack of comprehensive spectral libraries that are used to guide peptide identification and quantitative data extraction.

Here we offer a plasma proteomics spectral library in which we have utilized accessible tip-based fractionation and non-conventional proteases to boost proteome sequence coverage, and combined this with a DIA-MS strategy to reproducibly quantify the differential regulation of the plasma proteome upon active TB infection.

## Material and Methods

### Sample-specific library generation

Plasma samples from 3 adults with (0 with HIV) and 3 adults without (0 with HIV) active pulmonary TB were used from the FIND specimen bank. The samples were inactivated by addition of 2x inactivation buffer (8M urea, 100mM ammonium bicarbonate, 150 mM NaCl) in a 1:1 v:v ratio, followed by addition of RNAse (NEB) to 0.75*µ*L/mL concentration. 10 *µ*L of plasma from the individuals with active TB were pooled and depleted using the top12 most abundant depletion kit (Thermo-Fisher) according to manufacturer’s instructions. Following depletion, the samples were boiled at 90 °C for 5 minutes. Denatured proteins were reduced with 5 mM TCEP for 30 minutes at 56 °C and then alkylated with 10 mM of chloroacetamide for 30 minutes at room temperature in the dark. The samples were then loaded into a Vivaspin 3 KDa MWCO (Sartorius) and washed thrice with 200 *µ*L of MS-grade H_2_O. Samples were resuspended in 100 *µ*L of 50 mM ammonium bicarbonate and then subjected to proteolysis using either 2 *µ*g of trypsin (Promega), 2 *µ*g of AspN (Promega), or 2 *µ*g of GluC (Sigma-Aldrich) overnight at 37 °C on a shaker at 1000 rpm. Peptides were collected by centrifugation (8000 g for 30 minutes) and the filters were washed once with 100 *µ*L of ddH20. To perform basic reverse phase fractionation, the samples were acidified to 0.1% TFA final concentration. C18 spin columns (Nest group) were activated with 1 column volume of ACN and equilibrated with two column volumes of 0.1% TFA. Peptides were bound to the column and washed twice with 0.1% TFA. For elution, 7 solutions were used with increasing concentration of ACN in 0.1% triethylamine from 2.5% to 20% and following the last elution the column was washed twice with 1 column volume of 50% ACN (see Supplementary Table 1). Fractions were dried under vacuum and resuspended in 15 *µ*L buffer A (0.1% FA in MS-grade H20) and approximately 500 ng were subjected to proteomic analysis.

### In plate sample processing for Mtb positive and negative samples

5 *µ*L of plasma from individuals with and without TB diseases were inactivated following a similar procedure to the library generation and were separated into three samples (AspN, GluC, and Trypsin). Each samples was then loaded on a 96 well filter plate (Acroprep, PALL) with 3 KDa MWCO cutoff. Samples were washed twice with 200 *µ*L of MS-grade H_2_O. 50 *µ*L of TUA buffer (8M Urea, 5 mM TCEP, 25 mM ammonium bicarbonate) were addded and the samples were incubated on a thermo shaker at 37 °C and 400 rpm for 1 hour. Chloroacetamide was added to 10 mM final concentration and samples were incubated at room temperature in the dark for 30 minutes. Buffer was removed by centrifugation at 1000 RPM for 1 hr and samples were washed thrice with 200 *µ*L of MS-grade water and centrifuged to dryness. Proteins were resuspended in 50 *µ*L of 25 mM ammonium bicarbonate. 1 *µ*g of either trypsin, AspN, or GluC were added to each corresponding well and incubated on a shaker at 37 °C overnight. Peptides were recovered by centrifugation at 1000 rpm for 1 hour and plate was washed twice with 100 *µ*L of MS-grade H_2_O. Peptides were transferred to low-binding tubes and the receiver plate was washed with 100 *µ*L of 80% ACN to increase recovery of hydrophobic peptides. Peptides were dried under vacuum and resuspended in 12 *µ*L buffer A (0.1% FA in MS-grade H20). 3 *µ*L per tube was pooled together and pooled sample was defined as multi-enzyme digested sample (MS pool). Approx 500 ng were analyzed by mass spectrometry.

### DDA Pasef acquisition for spectral library generation

Data for each fraction was acquired on a timsTOF Pro mass spectrometer (Bruker) interfaced with a Thermo Easy-nLC 1200 (Thermo Fisher Scientific). The peptides were separated at a flow rate of 400 *nL/min* over a manually packed 15 cm long column containing 1.7 *µ*m BEH beads (Waters) packed with a silica PicoTip™ Emitter (inner diameter 75 *µ*m) (New Objective, Woburn, USA). Peptides were eluted from the column using a linear gradient from 2% to 32% buffer B (80% acetonitrile and 0.1% formic acid in HPLC grade H_2_O) in Buffer A (0.1% formic acid in HPLC grade H_2_O) with a total length of 90 minutes. The peptides were sprayed into the timsTOF Pro using a CaptiveSpray source (Bruker), with a end plate offset of 500 *V*, a dry temp of 200 °C, and with the capillary voltage fixed at 1.6 *kV*. The mass spectrometer was operated in positive ion mode. For DDA acquisition the timsTOF Pro (Bruker) was operated in PASEF mode using Compass Hystar v5.1 and oTOF control v6.2. The mass range was set between 100-1700 *m/z*, with 10 PASEF scans between 0.6 *V s/cm*^2^ and 1.6 *V s/cm*^2^. Accumulation time was set to 2 ms and ramp time was set to 100 ms. Fragmentation was triggered at 20,000 arbitrary units (a.u.) and peptides (up to charge 5) were fragmented using collisionally-induced dissociation (CID) with a spread between 20 *eV* and 59 *eV*.

### DIA Pasef Acquisition

For DIA acquisition, each sample was acquired on the same HPLC-MS setup previously described, and analyzed with either the 90 min gradient used for DDA analysis, or a shorter 50 minute gradient in which peptides were separated for 35 minutes using a linear gradient of buffer B (80 % acetonitrile and 0.1% formic acid in HPLC grade H_2_O) from 5% to 33%, then buffer B was increased to 40% in 5 minutes and the column was washed at 90% for 10 minutes before the next run. The separation was done at 400 *nL/min* while the column wash was performed at a flow rate of 500 *nL/min*. Similar MS1 range, PASEF parameters, and fragmentation parameters were employed as described above for DDA. 12 DIA-PASEF scans were performed.

### Mass spectrometry data analysis

#### Sample-specific library generation

The AspN library and trypsin libraries were generated using Spectronaut.^12^ The samples were searched using Pulsar against a combined database encompassing the *Mycobacterium Tubercolosis* proteome (4081 entries, downloaded from Uniprot on the 12/02/21) and *Homo Sapiens* proteome (20,397 entries, downloaded on 07/01/21). The default BGS settings without iRT normalization were used. The GluC spectral library was generated using MS-Fragger. ^13^ Briefly, the ‘SpecLib’ workflow was employed using default parameters. The number of missed cleavages was fixed to 2, using cysteine carbamydomethylation as fixed modification, N-terminal acetylation and methionine oxidation as variable modifications. The GluC DDA-PASEF files were also searched against the combined human-Mtb database. Decoys were generated by pseudo-inversion as previously described.^14^ Both searches were performed with 1% FDR at peptide and protein level. EasyPQP (https://github.com/grosenberger/easypqp,commit#dfa4ead) was used to generate the aligned retention time using high confidence iRT (ciRT). The resulting library was then converted into a Spectronaut-compatible library using an in-house Python script. The final sample specific spectral assay combined data from all proteases and encompasses 765,411 assays from 30,400 peptides mapping to 3309 protein groups (Supplementary Table 2). The spectral assay library has been deposited to the ProteomeXchange via the PRIDE^15^ partner repository with the dataset identifier PXD025671. To compute sequence coverage the protein coverage summarizer from the Pacific Northwest National Laboratory was used (https://github.com/PNNL-Comp-Mass-Spec/protein-coverage-summarizer).

#### Data processing and analysis for DDA and DIA data

DIA data for each protease was searched independently for both 90 minutes and 50 minutes gradients using Spectronaut and the correspondent spectral library. The settings employed in Spectronaut were default BGS (iRT normalization kit off) and each file was exported at the peptide level. For protein inference the average top3 peptide intensities were used. The resulting protein level matrix was log2-transformed and the data was normalized using median-centering. For missing value imputation, a distribution-based strategy was employed. For each sample, we selected the lowest 10% of values and calculated standard deviation (*σ*) and mean (*µ*). We then generated a normal distribution having similar *σ* but downshifted mean by 1.8 × *σ*. Rational for this imputation strategy is that lack of peptide detection cannot be differentiated between precursor ion intensity being below the limit of detection (LOD) or true biological absence. By sampling intensities below the LOD (defined here as the lowest 10% of recorded values per MS-injection) we assume that all not-detected peptides are below the LOD of the instrument. Following normalization and imputation the log2FC was calculated as ratios of the average intensities between Mtb infected and not infected individuals in log space. *P* were calculated using a two-tailed Welch t-test and corrected for multiple testing using the Benjamini-Hochberg correction. The coefficient of variation was calculated on the non-log transformed data and defined as 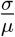.

For estimation of concentration for proteins detected in the spectral library, the concentration was downloaded from Human Protein Atlas^16^ (https://www.proteinatlas.org/humanproteome/). Concentrations were converted to *ng/L* and a quadratically penalized general linear model (GAM) was used for regression using logged intensity and logged concentration values. To estimate the concentration of Mtb proteins, the combined library was subset to only Mtb peptides and imported into Skyline v 20.2.0.343 (https://skyline.ms/project/home).^17^ Each transition was then exported for all spectral library DDA runs using its specific protease and fragment-level intensities. Transitions were summed up into peptides and then peptide intensities were average across proteases and fractions (in case of identification in multiple fractions) to obtain the overall protein intensity used for further concentration analysis. Interpolated value were used as is to estimate the concentration for the detected Mtb proteins and the rest of the identified proteins. Spectrums for ALKEGNER and DGRAVLR peptide were annotated using the IPSA tool.^18^ For enrichment analysis, Enrichr^19^ was used (https://maayanlab.cloud/Enrichr/) and the corrected *p* was used for all plots.

All data analysis was performed in python v3.8.1 (https://www.python.org) using pandas v1.1.3 (https://pandas.pydata.org), numpy v1.19.2 (https://numpy.org), ^20^ scikit-learn v0.23.2 (https://scikit-learn.org/stable/). ^21^

Figures 2 ACD, 3 AB, 4 AB, 5 ABCD and supplementary figures 2,3,4 were generated in R version 4.0.3 (https://www.r-project.org), using ggplot2 v3.3.2 (https://ggplot2.tidyverse.org) and ggpubr v0.4.0 (https://github.com/kassambara/ggpubr). Venn diagramms in figures 2B and supplementary figure 1 were generated using matplotlib v3.3.2 (https://matplotlib.org) and matplotlib-venn v0.11.6 (https://github.com/konstantint/matplotlib-venn). Figure 3 panel C and D were generated within the IPSA website. Workflow figure (Figure 1) was created using BioRender.com.

**Figure 1.**
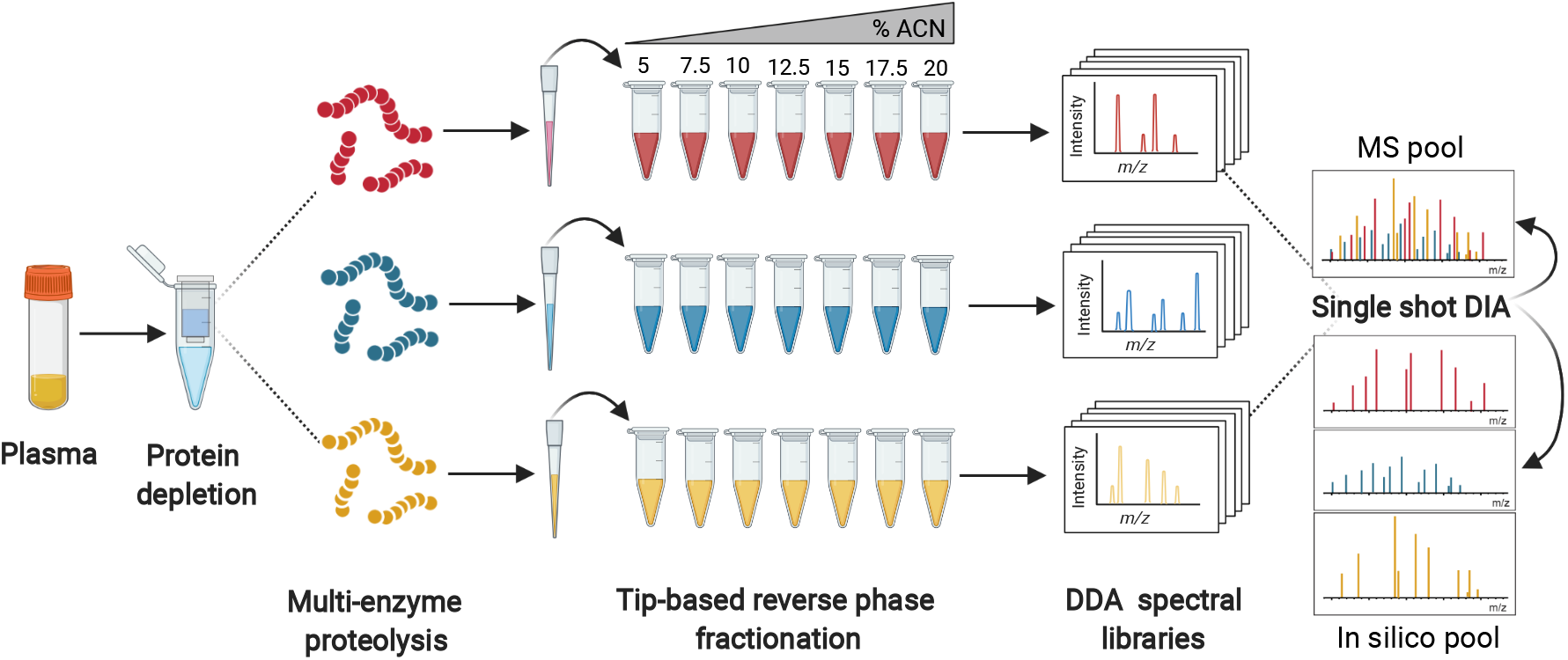
Schematic of the experimental workflow employed.

**Figure 2.**
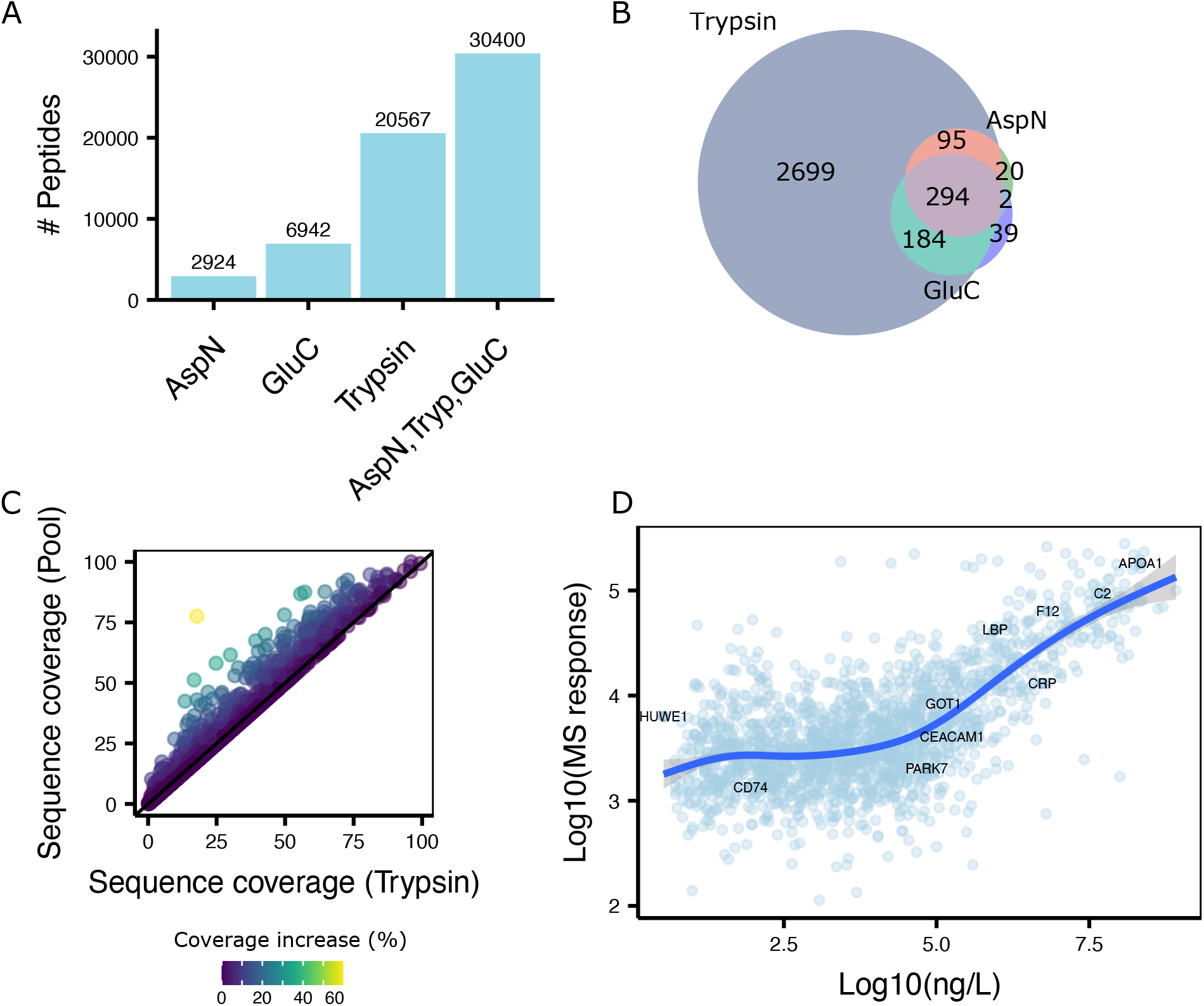
Description of the plasma spectral library derived from the combination of multiple proteases. **A** Barplot showing the number of peptides for each protease (AspN, GluC, Trypsin) and their combination. **B** Venn diagram showing the overlap of identified proteins for each protease. **C** 2D scatterplot illustrating the increase in sequence coverage by combined results from ApsN, GluC, and trypsin (Y axis) compared to only trypsin digestion (X axis). Each dots represents an individual protein. Color represents the percentage of increase in sequence coverage. **D** 2D scatterplot showing the estimated protein concentration from Human Protein Atlas^16^ on the X-axis and the MS response on the y axis.

**Figure 3.**
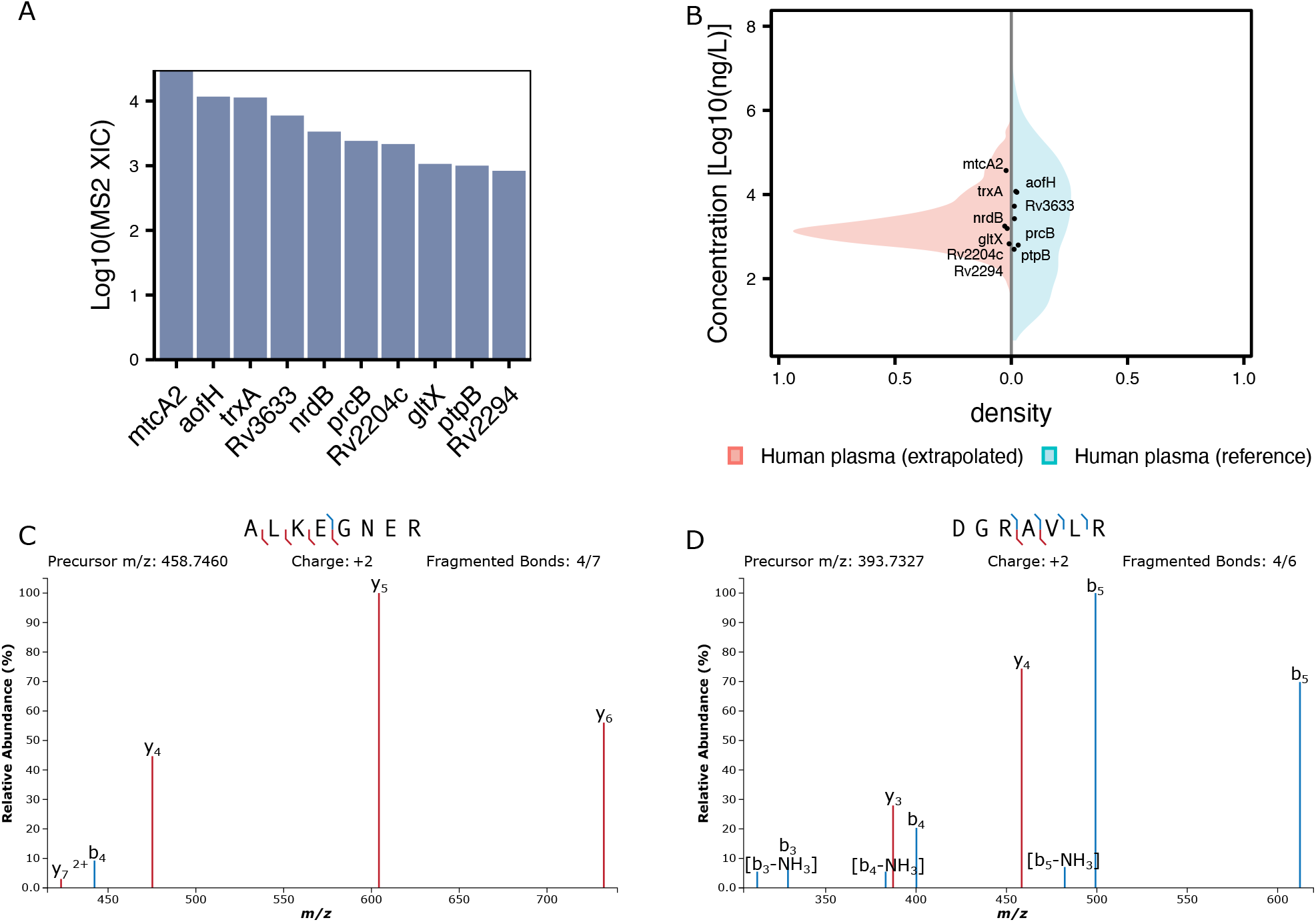
Coverage of Mtb proteins using orthogonal proteases. **A** Barplot showing recovery of Mtb proteins in the pooled spectral library. **B** Mirror plot showing the extrapolated concentration for TB proteins (black dots) versus the reference concentrations from the Human Protein Atlas (blue density) and the extrapolated intensity for the remaining proteins in the spectral library (red density). **C, D** Annotated MS2 spectrum for two proteotypic *mtcA2* peptides identified using Trypsin (C) or AspN (D).

**Figure 4.**
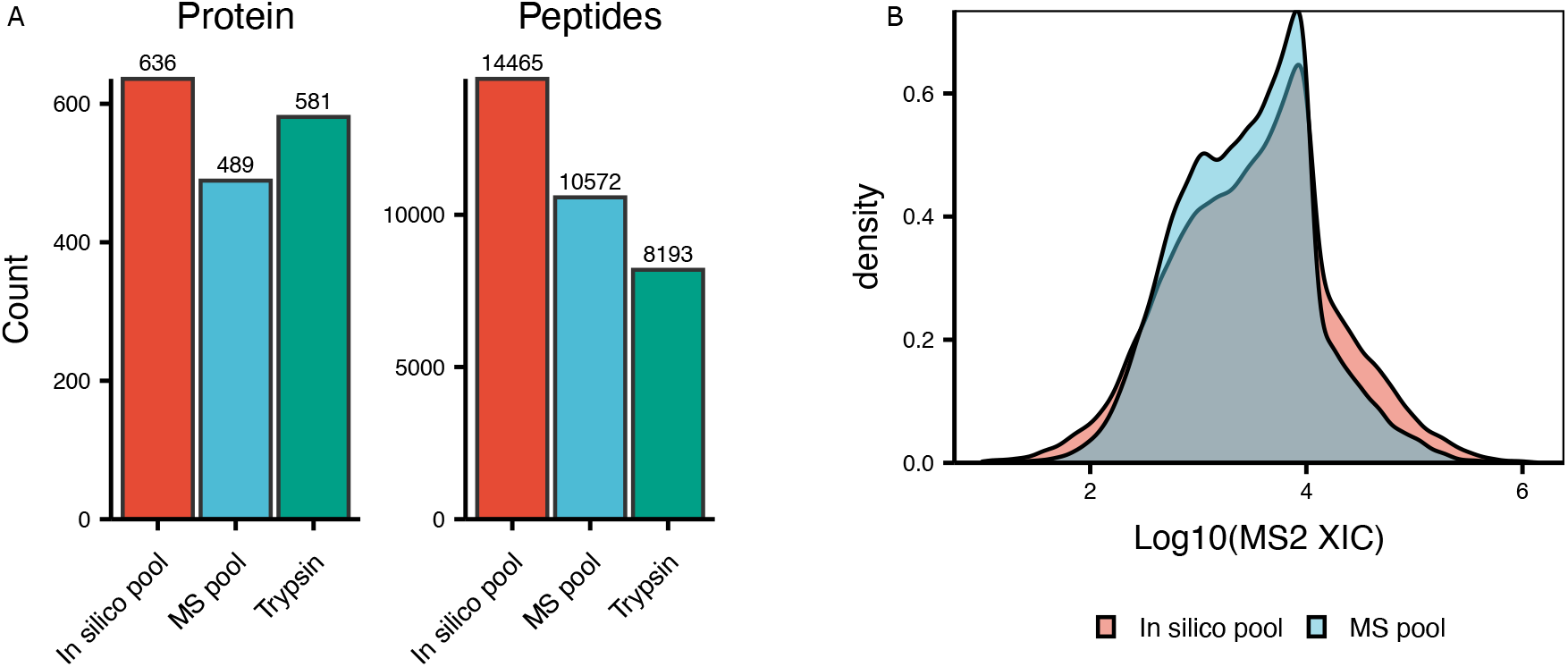
Comparison between *in-silico* pooled sample and MS acquired pooled sample. **A** Cumulative number of identified proteins and peptides from all proteases (in silico pooled sample), MS acquired pool and trypsin **B** Density plot for peptide intensities.

**Figure 5.**
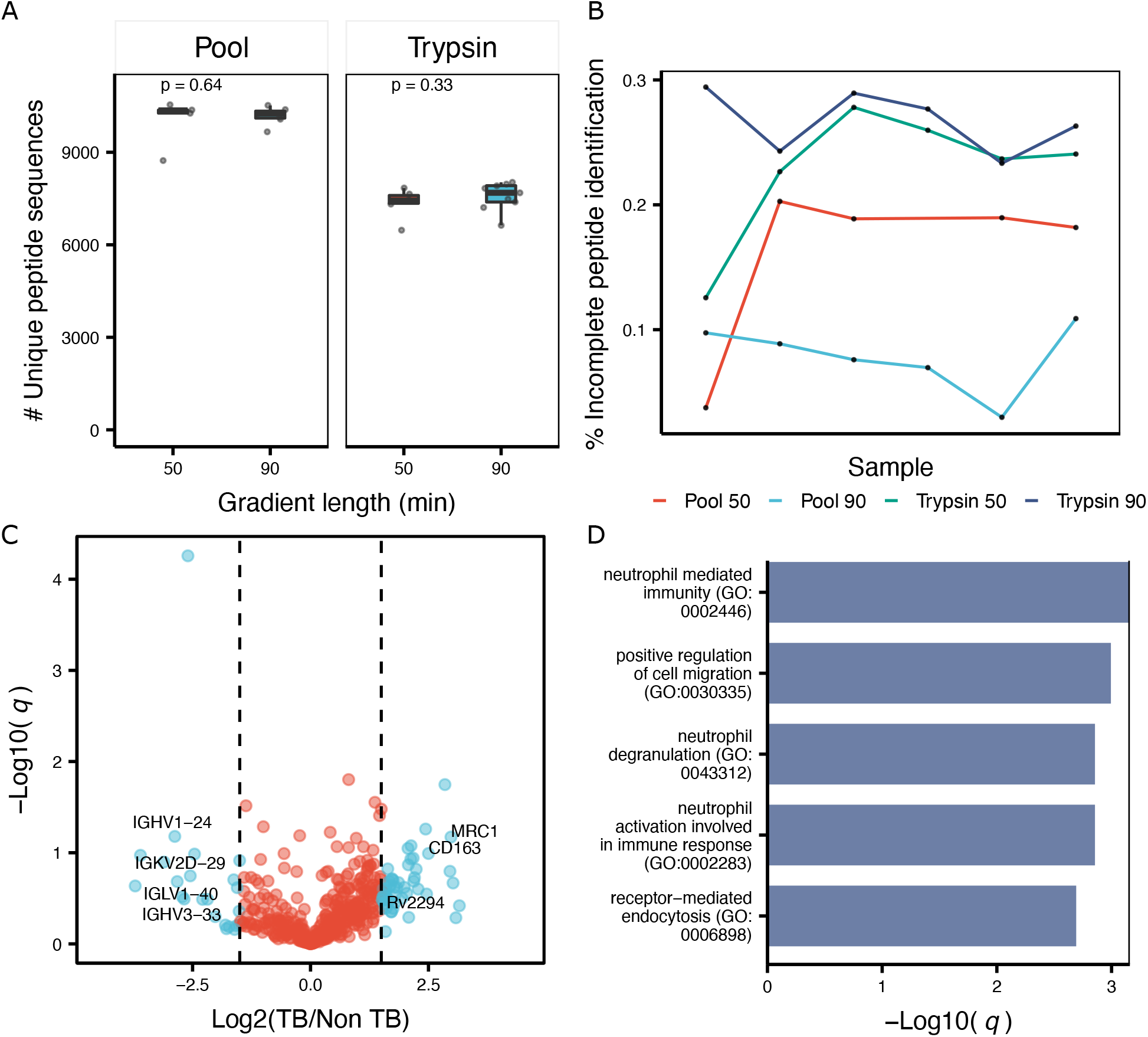
Differential analysis of Mtb infected samples using DIA-MS. **A**. Boxplot showing the number identified peptides by trypsin and the MS pooled sample using 50 and 90 minutes chromatographic gradients. Box represents the interquantile range (IQR) and its whiskers 1.5×IQR. Each dot represents one individual sample. P value represents the results of a paired Student t-test. **B**. Lineplot illustrating the number of missing values expressed as percentage of not consistently detected precursor ions using trypsin and the pooled sample. Color represents an enzyme and a specific gradient lenght (50 or 90) while samples are shown as black dots. **C**. Volcano plot for MS pooled data. X axis represent the log2 fold change. Y axis represents the negative log10 of the BH-adjusted p-value. **D**. Barplot showing the enriched GO terms for upregulated proteins. Bar represents the significance on the log scale from an hypergeometric test.

## Results

### Comprehensive plasma proteome spectral library generation

To reduce the sample complexity and facilitate the detection of low abundant proteins upon proteomic analysis, we first performed a depletion of high abundant proteins (Figure 1) and then individual aliquots of the depleted plasma were digested using either trypsin, AspN, or GluC. Finally, we applied a reversed-phase tip-based fractionation scheme (see method for details) under basic pH to generate orthogonal fractions and analyze each fraction in DDA-PASEF mode using a novel ion-mobility mass spectrometer. ^22^

The resulting spectral library encompassed in total unique 30,400 peptides of which 20,567 are derived from the trypsin digested samples, 2,924 from the AspN, and 6,942 from the GluC (Figure 2A). These numbers translate into 3,309 protein groups being identified across all proteases (Supplementary Table 2), with additional proteins being identified by digestion with either GluC or AspN (Figure 2B), possibly due to the generation of peptides more amenable for proteomics analysis.^23,24^ As expected, we observed an averaged increase in sequence coverage (7%) when combining AspN and GluC to the tryptic digested samples (Figure 2C). For 70% of the identified proteins we found annotation of their existence in plasma either in the Human Protein Atlas^16^ or Peptide Atlas,^25^ while 40% were found in both of these databases (Supplementary Figure 1). Notably, these databases are a combination of several hundred experiments, while we recapitulated a large portion of the identified proteins within a less than a day of MS acquisition.

Proteins were detected across 8 orders of magnitude based on their reported concentrations from the Secretome Atlas, ^16^ ranging from 3 *ng/L* (*HUWE1*) to *>* 8*e*^8^*ng/L* (*CP*); showcasing the great sensitivity of the TimsTOF Pro for detection of low abundant proteins (Figure 2D). We observed linearity between MS response and concentration (*R*^2^ = 0.88) over 5 orders of magnitude, suggesting a great degree of quantitative accuracy, which is essential for large scale biomarker studies.

### Identification and quantification of Mtb proteins

*Mycobacterium tuberculosis* (Mtb) proteins have been challenging to detect in plasma due to their intrinsic low abundance, estimated to be in the picomolar range,^26^ and their possible clearance by the immune system in immunocompetent individuals. In our samples from immunocompetent patients, we detected 10 Mtb proteins (Figure 3A) across all enzymes employed.

We proceeded to estimate the concentration of the Mtb proteins using generalized additive models (see Method for details). Among these proteins we identified those known to be secreted such as the tyrosine phosphatases *PtpB* ^27^ for which we observed one of the lowest estimated concentrations among all TB proteins detected (≈ 6−7*µg/L*). Interestingly, while the proteins expressed at the highest abundance in the infection site (lungs) are reported to be the component of the cholesterol metabolism and nitrogen processing pathways,^28^ we identified additional metabolic enzymes such as *nrdB* and *mtcA2*, potentially suggesting these proteins are secreted or more likely released after clearance of Mtb by immune cells. To further support the presence of *mtcA2* in the analyzed plasma sample, we manually extracted all identified peptides using Skyline^17^ for all proteases employed (Figure 3C and D). Comprehensive fragment coverage and the presence of two proteotypic peptides for this protein are observed, confirming its presence in our samples. Among the other proteins detected, we observed the transporter *Rv2994* which is an uncharacterized Mtb protein recently shown to be clinically valuable for Mtb serodiagnosis.^29^ Lastly, *Rv2204c* has also been shown to be a marker of active and latent Mtb infection.^30,31^

### DIA analysis of multi-protease digested Mtb infected samples

DIA analysis of each protease sample individually resulted in the combined detection (*in silico* pool of 14,665 peptides (636 proteins), the majority of which resulted from trypsin digestion (Figure 4A). Each sample was analyzed also both using a short (50 minutes) or a longer (90 min) chromatographic gradient. We found the number of proteins or peptide did not significantly increase with longer gradients (Figure 5A, Supplementary Figure 2), highlighting the fast duty cycle of qTOF mass spectrometers.^32^

We also mixed samples from each protease into a single samples and performed DIA of this pooled samples (MS pool). Comparison of the MS-pooled sample to the *in-silico* generated one showed a recovery of 73% (10,572/14,465) at the peptide level compared to the *in-silico* pooled sample (as depicted in Figure 4A) albeit at a reduced number of proteins identified (489) compared to trypsin (581). We hypothesized this effect was dependent on the presence of several high-abundant peptides in each protease-digested sample, and that such pooling masked the detection of low abundance peptides due to an increase in the total fraction of the sample comprised of high abundance peptides. Indeed, when comparing the distribution of detected peptides in the MS-pooled sample and the *in-silico* pooled sample, we observed a decrease in identification of low-abundant peptide precursors in MS-analyzed sample, with an increase in high-abundant peptides (Figure 4B) corroborating our hypothesis. Additionally, the MS-pooled DIA data showed great consistency of protein identification (Figure 5B) and quantitation resulting in only 17% of incomplete features (defined here as peptides not consistently identified across all samples), which outperforms the trypsin digested samples by ≈ 6% in the 50 minutes gradient and ≈ 20% in the 90 minutes gradient. This consistency resulted in an average coefficient of variation (CV) of 38% (Supplementary Figure 3A) for the pooled data, approximately 8% less of the tryptic samples (*p* = 1.5×10^*−*5^) and an overall lower number of missing values across samples (Supplementary Figure 3B). While statistically underpowered due the small number of samples analyzed, we identified 34 proteins being enriched in the TB diagnosed samples compared to the control samples. Among the top dysregulated proteins we found several proteins which are known to be involved in TB pathology. For example, we observed elevated (Log2 fc = 2.97) Macrophage mannose receptor 1 (*MRC1*) protein which is a C-type lecitin responsible for recognition of bacterial infection.^33^ Additionally, we found the protein cluster of differentiation 163 (*CD163*) increased upon active TB. This protein mediates the transition from monocyte to macrophages and has been previously reported to be of clinical relevance as a biomarker of treatment efficiency and overall diseases progression. ^34^ Unsurprisingly, the majority of the upregulated proteins are part of inflammatory pathways (Figure 5D) which shows the burden of the immune system in Mtb infected individuals. Gene-disease association analysis revealed the enriched proteins to be primarily associated with pneumonia (Supplementary Figure 4). When analyzing the downregulated proteins, we observed several immunoglobulins having lower abundance in our TB cohort compared to the healthy controls. Interestingly, this has also been observed in another proteomics study.^7^ Overall our analysis recapitulates previous findings and showcases the applicability of DIA and multi-protease digestion for robust analysis of clinical samples.

## Discussion

Clinical proteomics play an important role in understanding the pathogenesis of human disease and identifying new biomarkers for diagnosis and treatment monitoring. As plasma is easy to obtain and commonly used in diagnostic testing, we developed a novel protocol that utilizes orthogonal proteases coupled with DIA-MS to improve dynamic range, protein coverage, and quantification. While mass spectrometry has not been routinely used in large scale clinical trials and biomarker discovery cohorts, it has the potential to be a key technology for robust protein detection and quantification in a variety of clinical settings. We have demonstrated its utility in TB disease, which triggers a large host response and creates a complex plasma sample that can challenge standard mass spectrometry approaches.

From a biological perspective, our results recapitulate several previous transcriptomic and proteomic analyses from TB patient samples, such as the upregulation in inflammatory pathway components reported to be specific for TB disease.^35^ The sensitivity of our methods enabled the recovery of nearly half of the previously reported plasma proteins within a single fractionation experiment and resulted in the identification and quantification of diagnostic Mtb proteins in plasma, which were previously accessible primarily by antibody-based assays. Interestingly, this included the detection of intracellular Mtb proteins not expected to be secreted and suggests the intriguing hypothesis that, even before treatment, a fraction of Mtb is cleared and the proteins are released in the circulation. The recent discovery of several mechanisms by which the pathogen releases extracellular vesicles^36^(EVs) could also provide an explanation for our observation. While none of the Mtb proteins detected here have been reported in Mtb vesicles,^37^ EVs composition is known to vary^38^ thereby more work is needed to highlight the compositional heterogeneity of Mtb vesicles. Thus, these findings highlight the need for unbiased analysis of biofluids to gain insights into TB biology.

The rapid development of DIA-MS shows great potential towards biofluid analysis, however previous studies were limited in the number of proteins identified due the lack of comprehensive spectral assay libraries. Here we shown the use of non-conventional proteases combined with DIA-MS to increase coverage of the plasma proteome. The combination of multiple proteases within a single sample improved identification and quantification robustness, which are key features for technologies currently applied in modern diagnostic (PCR, NGS, etc). While we observed a slight decrease in protein identifications upon pooling proteases in DIA analysis, the proteins additionally identified by trypsin were not consistently found across samples and are thus unlikely to have potential clinical utility.

Altogether, we showcase the applicability of library-based DIA-MS for plasma proteomics for consistent recovery of hundreds of proteins with a great degree of quantitative accuracy. We anticipate our spectral library can serve as a useful as a base for future biomarker studies utilizing the timsTOF Pro, or complemented with additional assays to increase proteome coverage. While our approach showed improvements over previous methods, a limitation is that current tools for DIA analysis, and more broadly DIA acquisition, have been developed specifically for tryptic digests. Thereby it is conceivable to develop ad-hoc DIA windows schemes which exploit differences between proteases (e.g. *z, m/z*, etc.) to more comprehensively sample the precursor space while reaching an optimal duty cycle. Further advances in software could also include FDR models trained on non-tryptic sets or novel decoy-generation methods may also significantly improve the number of peptides which are possible to extract from DIA data using alternatives proteases. Looking forward, the application of alternative proteases could be beneficial to perform deep proteomic profiling of clinical specimen and to increase the confidence in identified proteins in large clinical cohorts.

## Conclusions

We used digested plasma from different proteases and acquired them in DIA-MS using a library derived from a tip-based fractionated representative plasma sample. We showed increased sequence coverage, robustness, and reduced missing values for the combination of AspN, GluC, and trypsin compared to a standalone tryptic digested sample.

## Acknowledgements

We thank Nevan J. Krogan for use of the Thermo Fisher Scientific Proteomics Facility for Disease Target Discovery at the Gladstone Institutes. We thank FIND for providing plasma samples from its specimen bank.

## Funding

NIH R01GM133981 to DLS, NIH R01AI152161 to AC and JE, and NIH K23HL153581 to DJ.

## Author contributions

AF, DLS, ALR conceived and designed the project. AF and KC performed the experiments. AF and ALR performed the data analysis, AF, DLS, ALR drafted the manuscript. All authors critically reviewed the manuscript and approved the final version. DLS supervised the work.

## Competing interests

None.

## Data and materials availability

All raw MS data files, search results and individual spectral libraries are available from the Pride partner ProteomeXchange repository under the PXD025671 identifier with username: and Password: OZfwGGiA.

## Supplementary figures

**Supplementary Figure 1.**
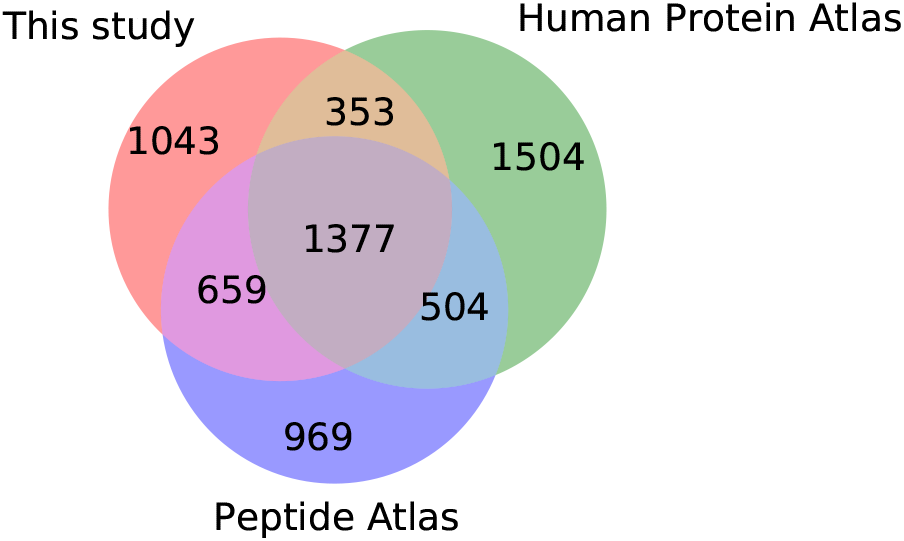
Overlap between gene products identified in this study and other plasma protein databases.

**Supplementary Figure 2.**
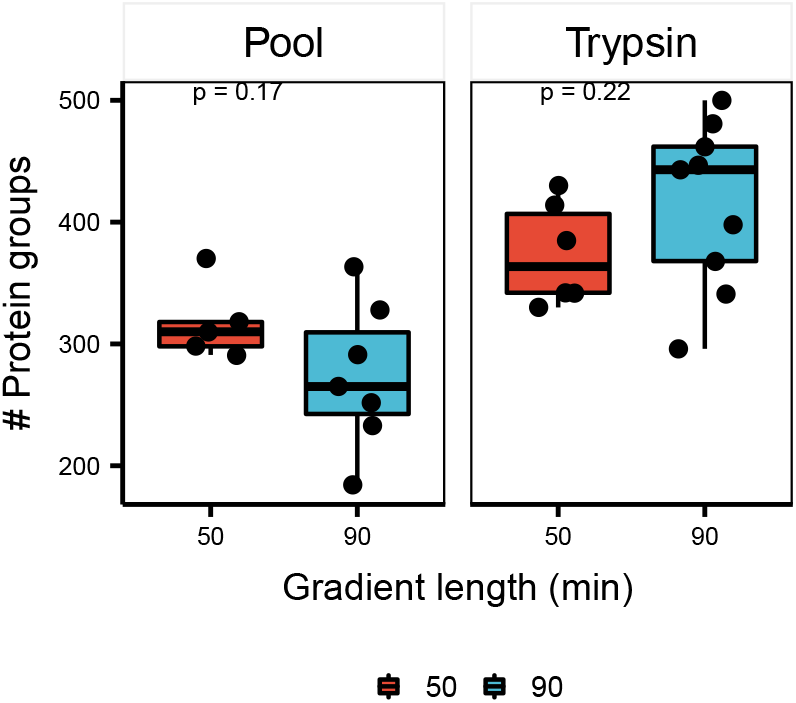
Protein identification comparing long and short gradients. Boxplot showing the number of identified proteins at 1% FDR using either 50 minutes or 90 minutes chromatographic gradient. The box represents the IQR and its whiskers 1.5×IQR, while the black line highlights the mean. Individual samples for each protease are reported as dots.

**Supplementary Figure 3.**
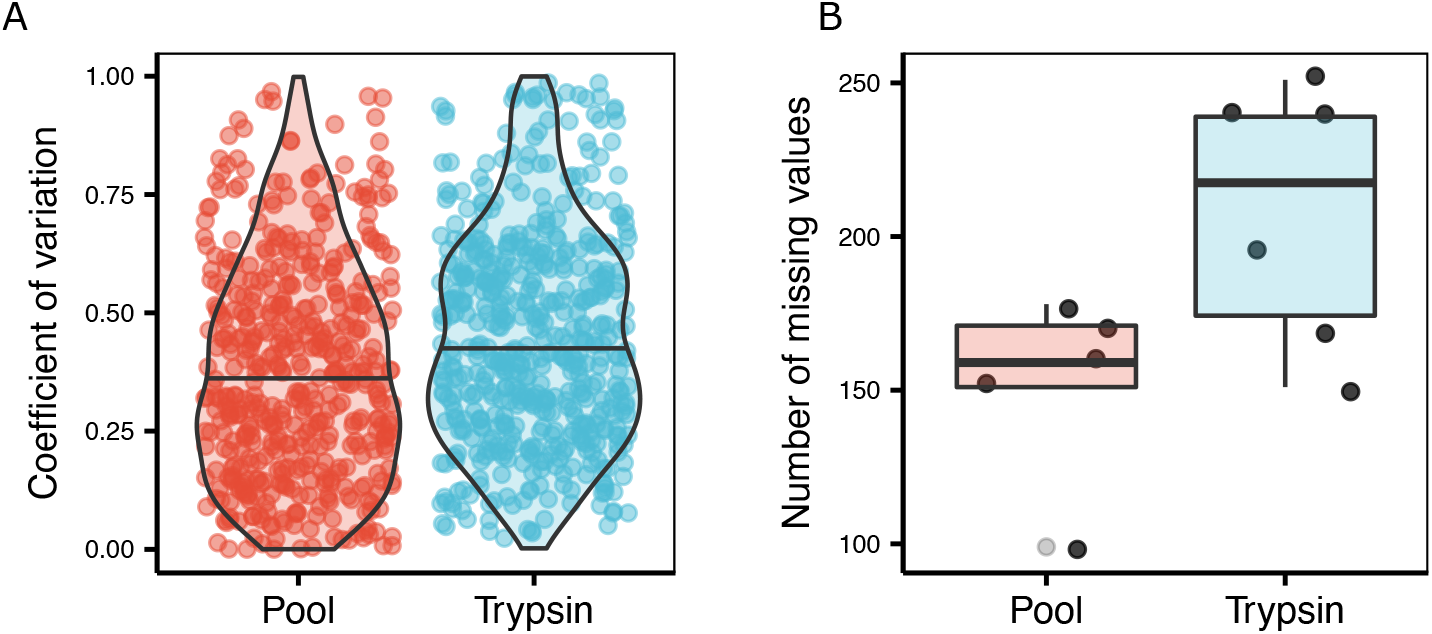
Consistency of quantification and identification for pooled DIA data using 50 minutes chromatographic gradient. **A** Violin plot representing the coefficient of variation at the protein level for the DIA data. Each dot represents a protein **B** Boxplot showing the missing value counts per sample. The box represents the IQR and its whiskers 1.5 × IQR, while the black line highlights the mean. Individual samples for each protease are reported as dots.

**Supplementary Figure 4.**
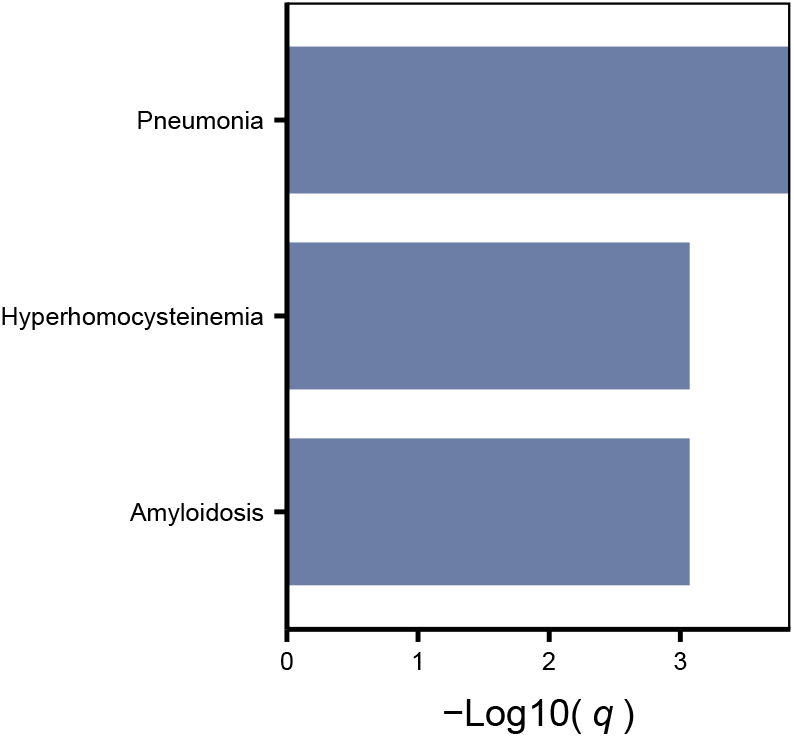
Barplot for enriched disease terms from the DISEASES database. Bar represents the −*log*10 of a p-value derived from an hypergeometric test. The Bonferroni procedure was applied for multiple testing correction.

## Supplementary tables

**Table 1.** Reverse phase fractionation scheme employed.

| Fraction number | ACN% |
| --- | --- |
| 1 | 5% |
| 2 | 7.5% |
| 3 | 10% |
| 4 | 12.5% |
| 5 | 15% |
| 6 | 17.5% |
| 7 | 20% |
| 8 | 50% |

**Table 2.** Description of spectral library derived from AspN, GluC and Trypsin digested plasma.

| Enzyme | Assays | Peptides | Protein groups |
| --- | --- | --- | --- |
| AspN | 69557 | 2924 | 411 |
| GluC | 155787 | 6942 | 519 |
| Trypsin | 540657 | 20567 | 3272 |
| Combined | 765411 | 30400 | 3309 |

## Notes

### Competing Interest Statement

The authors have declared no competing interest.

